# The effect of removing repeat-induced overlaps in *de novo* assembly

**DOI:** 10.1101/2023.04.16.537101

**Authors:** Ramin Shirali Hossein Zade, Thomas Abeel

## Abstract

Determining accurate genotypes is important for associating phenotypes to genotypes. *De novo* genome assembly is a critical step to determine the complete genotype for species for which no reference exists yet. The main challenge of *de novo* eukaryote genome assembly, particularly plant genomes, are repetitive DNA sequences within their genomes. The introduction of third generation sequencing and corresponding long reads has promised to resolve repeat-related problems. While there have been notable improvements, reads originating from these repeats are still creating errors because they introduce false overlaps in the assembly graph. This study focuses on analyzing the effect of repeats on *de novo* assembly and improving performance of existing *de novo* assembly algorithms by removing repeat-induced overlaps. First, we show the possible improvements in de novo assembly with removing repeat-induced overlaps. Then we propose several methods for detecting and removing repeat-induced overlaps and evaluate their performance on several simulated datasets.

## Introduction

The goal of *de novo* genome assembly is to reconstruct a species’ genome sequence as completely as possible using a large number of relatively short sequences referred to as “reads” that are read from the species’ genome. While high-quality assemblies are already available for many species, many branches of the tree of life still need representative genome sequences. Recently, due to the popularity of long-read sequencing technologies, *de novo* assembly has once more become of interest. In this paper, we focus on improving the standard long read *de novo* assembly pipeline.

Most *de novo* assembly pipelines suitable for long reads follow the OLC paradigm: overlap-layout-consensus. First, in the overlap step, pairwise alignments between the reads are identified. The output of the overlap step is a set of pairwise read overlaps that can be represented as a graph, where nodes are the reads, and edges indicate overlaps between the reads. This graph will be referred to as the assembly graph. Second, the layout step tries to identify bundles of overlaps that belong together. This is done by pruning unwanted edges from the graph such that it becomes more linear through several graph cleaning procedures. Once all procedures are done, the graph is split up into contigs. Finally, the consensus step of the assembly pipeline identifies the most likely base for each position. The layout step is arguably the most differentiating step between the various *de novo* assembly methods that exist. This can go from extremely simple, e.g. miniasm (1) to very intricate with many manually optimized rules and corresponding specific data types, e.g. DISCOvar (2).

A problem that has plagued *de novo* assembly since the beginning is interspersed repeats in the species’ genome sequence. The interspersed repeats are sufficiently similar sequences that occur in two or more distinct genomic locations. The reads originated from any of the repeat instances introduce pair-wise overlaps with all instances of the repetition across the genome, which leads to cross-connections in the assembly graph. This will confuse the ‘layout’ step in the OLC assembly paradigm. Reads spanning the repetitive region can resolve the confusion by connecting the two sides of the repetitive regions together. While read lengths have been increasing dramatically for Third Generation Technologies (TGS), for the vast majority of eukaryotic species, the read length is still orders of magnitude smaller than the genome size. Moreover, it is unlikely that we will experience the luxury of chromosome-spanning reads like the ones observed for some microbial genomes soon (3–5). Finally, TGS reads are often still not (yet) long enough to span most of the repetitive regions in eukaryotic genomes.

In this paper, we analyze the effect of interspersed repeats on *de novo* assembly. Next, we show that removing repeat-induced overlaps can improve the performance of *de novo* assembly in different eukaryotic genomes, e.g. yeast, human, and potato. We demonstrate that a perfect classifier can increase the coverage of genome assembly by 0.1%, 4% and 7% in yeast, potato, and human chromosome 9, respectively. Finally, we also investigate some methods to detect and remove repeat-induced overlaps and compare their performance to the standard *de novo* assembly pipeline. Initially, we tried a baseline method and removed overlaps based on their degree in the assembly graph. Second, we trained a machine-learning model to detect and remove repeat-induced overlaps based on GraphSage node embeddings (6). While this method makes the overlaps set much smaller, it is not improving the assembly performance and the results are close to the standard *de novo* assembly pipeline.

## Material and methods

### Data

#### Reference sequences

In this study, we use the reference sequences of three species with differing degrees of repetitive sequences: *S. cerevisiae* (yeast) and *S. tuberosum* (potato), and *H. sapiens* (human) chromosome 9, which is the most repetitive chromosome in the human genome. We use high-quality available reference sequences as the source to simulate reads. We retrieve sequences from Genbank: yeast S288C genome assembly R63 (GCA_000146045.2), potato DM_1-3_516_R44 genome assembly version 6.1 (GCA_000226075.1), and human genome assembly T2T-CHM13v2.0 (GCA_009914755.3).

The potato reference sequence contains Ns to fill the gaps and unplaced sequences, complicating analysis. The Ns make problems for the evaluation step because we need a complete genome to compare the assemblies with it. We remove the unplaced sequences and the Ns to make the experiments straightforward. After removing Ns and unplaced contigs, we have one complete sequence for each chromosome.

#### Detecting interspersed repeats

We use Generic Repeat Finder (7) version 1.0 with the default parameters to detect interspersed repeats in these three reference sequences.

#### Simulating reads and genomes

We use aneusim (8) version 0.4.1 with default parameters to simulate diploid sequences (ploidy=2) close the reference sequences but with mutations and translocations. We use the simulated haplotype 1 and 2 sequences as genomes of two other individuals of these organisms for further analysis.

We use SimLoRD (9) version 1.0.2 to simulate reads similar to PacBio with 40x of coverage (-c 40) from the reference, and the simulated sequences. Using simulated reads allows us to label the alignments between the reads since we know where the reads originated from.

#### Alignments and labeling

We use minimap2 (10) version 2.13-r858-dirty with the default parameters to find the pairwise alignments between the reads. We label each alignment according to the origination coordinates of the reads participating in it. If the origination coordinates of the reads participating in an alignment overlap, then we label the alignment as a normal overlap. Otherwise, we label the overlap as a repeat-induced overlap.

### Genome assembly and evaluation

We use the miniasm (1) version 0.3-r179 with default parameters to assemble the sets of overlaps before and after intervening and removing the candidate alignments.

We use compass (11,12) to evaluate the de novo assemblies. While compass reports many metrics, we only report coverage, validity, multiplicity, the number of contigs and the longest contig. Supplementary Table 1 list the metrics and explain them. Coverage is the most important metric for this study because it shows what percentage of the genome is missing in the assemblies and can show us how much extra sequence, we achieve by removing repeat-induced overlaps. Another important metric is the number of contigs representing the assembly’s contiguity. It is essential to achieve higher coverage while maintaining the contiguity of the assembly.

### Feature extraction and training classifier

We use the reference sequences and the first simulated haplotypes as the training set and the second simulated haplotypes for the test. To train the model, first, we need to extract features for each overlap based on the assembly graph.

First, we create the graph using networkx (13) version 2.8.4. Then, we train a GraphSage (6) model on the assembly graph using the StellarGraph (14) library version 1.2.1 while the only attribute we add to the nodes is their degree. To learn the embeddings, we make a model which gets two nodes as input and predicts if there is a normal edge, repeat-induced edge, or no edge between them. Our model consists of three GraphSage layers with followed by a softmax layer for the prediction. We use categorical cross entropy as the loss function and Adam optimizer to train the network (learning rate = 0.001). This model contains 3 GraphSage blocks, which each contains 50, 50, 20 GraphSage layers, respectively. Moreover, the network iterates each GraphSage block 20 times before delivering the output to the next block. We train the network for 20 epochs and the batch size is 50. Since GraphSage models are inductive, after training the model, we can use the output of GraphSage layers to get the node embeddings in other graphs.

However, because the assembly graphs are huge, we need to subsample the graph for training and testing the model. We use the edgesampler module in the StellarGraph library to get the subgraphs. For yeast sequences, we take 20% of the nodes for training and 20% of the nodes for testing, while for human sequences, we use 2% of the nodes for training and 2% for testing.

Then, we use GraphSage embeddings to train a logistic regression classifier for separating repeat-induced and normal overlaps. We use the first simulated dataset to train this classifier. First, we create the assembly graph of the simulated dataset, and then extract the node embeddings using the previously trained GraphSage.

We use the GraphSage model to extract node embedding for every node in the assembly graph, and we concatenate embeddings of the two nodes participating in an edge, to get embedding of that edge, which represents an overlap. After creating the embedding of each overlap, we use sklearn (15) version 1.0.2 to train a logistic regression classifier with parameter C=0.001 to detect repeat-induced overlaps. We use 10-fold cross-validation to evaluate the classifier and select the model with the highest F1 score.

Finally, we use the GraphSage model to extract the embeddings of the second simulated dataset. Then we use the selected model from the previous step to remove overlaps classified as repeat-induced. Next, use miniasm (1) version 0.3-r179 to assemble the remaining overlap set and compare the results with the standard genome assembly pipeline.

## Results and discussions

### Characteristics of interspersed repeats in yeast, potato, and human genomes

In the first step, we used Generic Repeat Finder to detect interspersed repeats in the genome of yeast, potato, and human chromosome 9. Table 1 shows the statistics of the interspersed repeats available in these genomes. There are gaps in the potato reference sequence, which are indicated by Ns in the sequence. To simplify the analysis, we removed Ns from the reference sequence. Unresolved repeats are usually responsible for most Ns in the sequence. Consequently, in Table 1, we report fewer interspersed repeats for the potato genome than are present. The analysis is also simplified for human chromosome 9 since it is separated from the rest of the chromosomes, thereby excluding the occurrence if interspersed repeats in the other chromosomes from the analysis.

**Table 1:** The amount of interspersed repeats in yeast, potato and human chromosome 9 genomes.

| Organism | Genome size | Number of repeats | Repeat content (%) |
| --- | --- | --- | --- |
| Yeast | 12Mbp | 4022 | 28Kbp (0.2%) |
| Potato | 731Mbp | 8582087 | 76Mbp (10.3%) |
| Human chromosome 9 | 150Mbp | 625288 | 9Mbp (6%) |

As shown in Table 1, the repeat content is much higher in human chromosome 9 and potato than in yeast. Around 10% of a potato genome is interspersed repeats, which shows the high repetitive content in that is a hallmark of plant genomes. Human chromosome 9 contains 6% interspersed repeats, but this number may be higher if the entire genome is considered. There are only 0.2% interspersed repeats in yeast’s reference genome, indicating a simpler genome architecture.

The distribution of interspersed repeats follows a similar pattern in the three test organisms. However, human chromosome 9 has many longer repeats than the other two organisms (see Figure 1). As mentioned before, the count of repeats in the human genome can be even more than what is shown in Figure 1 because they might also be present in other chromosomes, which we did not consider in this study. Interestingly, although yeast has lower repeat content (see Table 1) than the other two organisms, it has some very long repeats. The longest repeats in the yeast genome are even longer than the potato’s longest repeats. However, this is likely due to the fact that the potato reference sequence is incomplete and the Ns are representing unresolved repeats.

**Figure 1:**
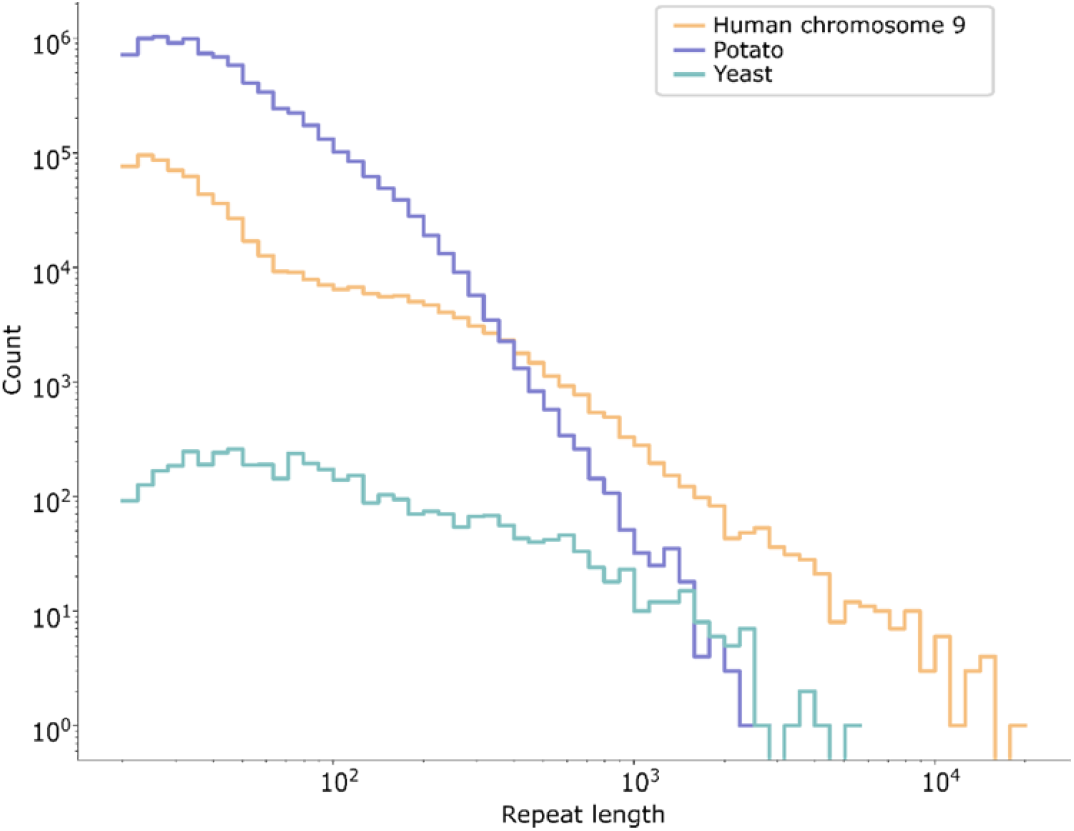
Histogram of the length distributions of interspersed repeats on chromosomes 9, potato, and yeast. In these three organisms, most interspersed repeats are smaller than 1000 bp. Despite this, all three organisms have repeats longer than 1000 bp, which complicates the *de novo* assembly process, as not all long reads will span the repeats completely.

The number of times each repeat occurs varies from 2 to more than 1000 times in the three model organism (see Figure 2). There are interspersed repeats in Human chromosome 9 that occur more than 40000 times, without considering other chromosomes that these repeats might be present. It is worth noting that the smaller repeats occur more often through the genome (see Supplementary Figure 1).

**Figure 2:**
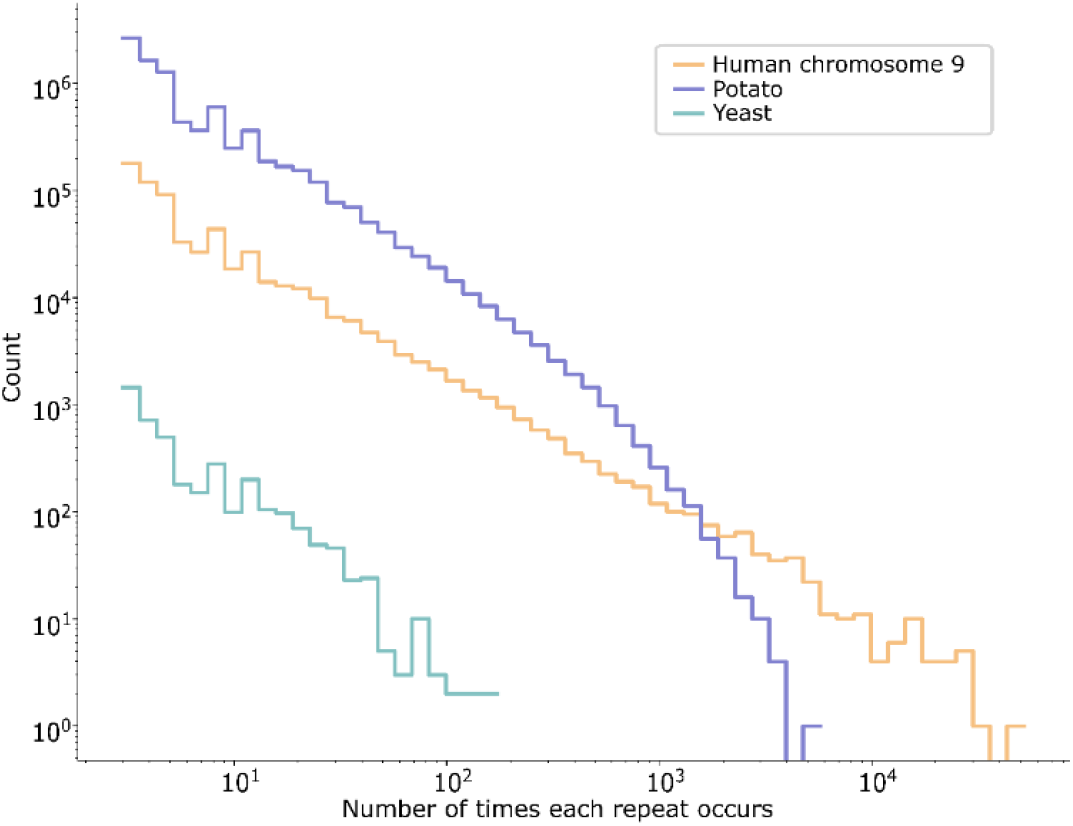
Histogram of the number of times each repeat occurs in the genome. The majority of interspersed repeats occur less than 100 times, but there are repeats in potato and human genomes that occur more than 1000 and 10,000 times, respectively.

### The effect of interspersed repeats in genome assembly

Next, we inspected the effect of interspersed repeats in genome assembly based on simulated reads from the reference genomes. Since the simulator reports the coordinates where a simulated read originated from, it is possible to label the pairwise alignment of reads. If there is an alignment between two reads but the coordinates these reads are sampled from do not overlap, we considered the alignment as repeat-induced. Otherwise, we labeled the alignment as normal. Table 2 shows the number of repeat-induced edges in yeast, human chromosome 9, and potato.

**Table 2:** This table shows the number of repeat-induced and normal edges in the assembly graphs. Although human and potato have only 6% and 10% repetitive sequences in their genomes, they have 71% and 96% repeat-induced edges in their assembly graphs.

| Organism | Repeat-induced edges (%) | Normal edges (%) |
| --- | --- | --- |
| Yeast | 189842 (8%) | 2093297 (92%) |
| Potato | 308658703 (96%) | 12084513 (4%) |
| Human chromosome 9 | 63004592 (71%) | 25221954 (29%) |

Reads that originate from one of the interspersed repeats align with reads from all other instances, which creates repeat-induced edges in the assembly graph. The human and potato reference sequences have considerably high repetitive sequences. Therefore, in the human and potato assembly graphs, the majority of the edges are repeat-induced in their assembly graphs (see Table 2). Subsequently, the reads originating from interspersed repeat regions also have a high degree in the assembly graph. Figure 3 shows the degree of the normal and repeat-induced edges in the assembly graphs. We define the degree of an edge as the sum of the degree of the two nodes connected by the edge. Figure 3 shows that most edges with a degree greater than 1000 are repeat-induced.

**Figure 3:**
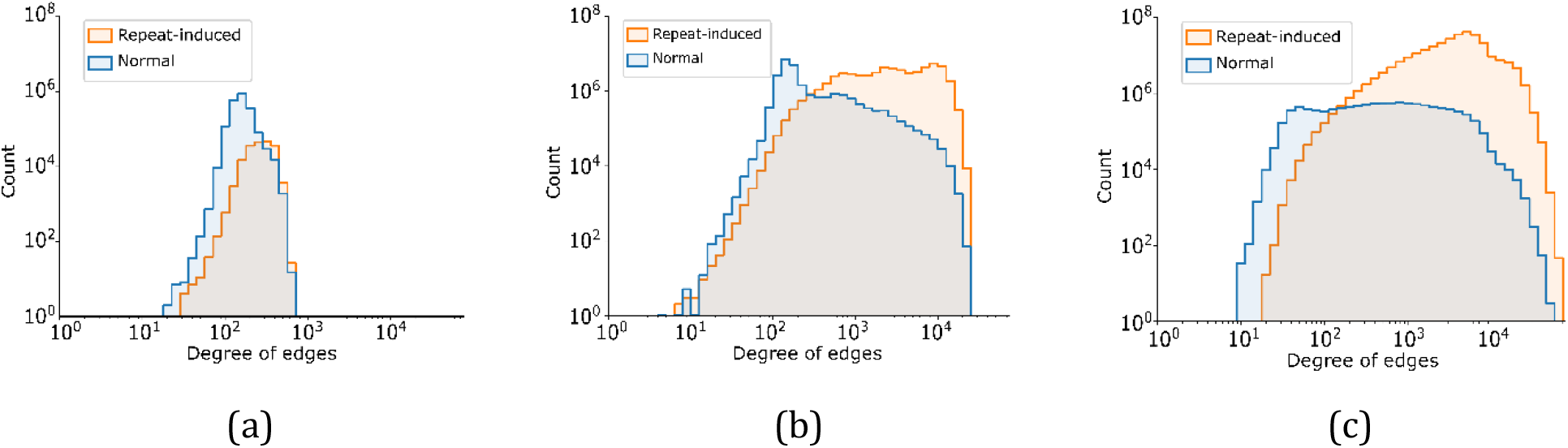
This figure compares the histogram of the degrees of repeat-induced and normal edges in the assembly graphs of yeast (a), human chromosome 9 (b), and potato (c). The degree of an edge is defined as the sum of the degrees of the two nodes it connects. There is no significant difference between the degree of repeat-induced and normal edges in the yeast assembly graph. On the other hand, in human chromosome 9 and potato, most edges with degrees greater than 1000 are repeat-induced.

To analyze the effect of repeat-induced overlaps in the assembly, we evaluated assemblies in the three model organisms before and after removing repeat-induced overlaps. In the normal scenario, we aligned the reads with minimap2 and assembled the genome with miniasm, reads, and the overlaps from the last step. In the removing repeat-induced overlaps scenario, we intervened in the assembly process, removed all the alignments labeled as repeat-induced, and used miniasm to assemble the remaining overlaps set. Table 4 shows the results of these two scenarios in the three model organisms. In all three datasets, removing repeat-induced overlaps improves genome assembly. In the yeast genome, removing repeat-induced overlaps lead to 6% more coverage. In the potato genome removing repeat-induced overlaps lead to 8% more coverage. This is expected since the potato genome is much more repetitive than yeast and suffers from more repeat-induced edges. In the human chromosome 9 dataset removing repeat-induced edges lead to 3% more coverage.

We tested whether removing a percentage of repeat-induced overlaps would still improve assembly performance in another experiment, where we removed 25%, 50%, and 75% of repeat-induced overlaps in the human chr9 genome and compared the final assemblies. It is clear from Table 3 that removing more repeat-induced overlaps improves coverage and validity and increases the length of the longest contig. However, the multiplicity, number of contigs and the assembly size is increasing after removing 25%, 50%, 75% repeat-induced overlaps and finally drops and get closer to one after removing all of the repeat-induced overlaps. This means by removing a portion of repeat-induced overlaps the assembler is replicating some of the repetitive regions which are valid sequences, but increases multiplicity and assembly size. Finally, with removing all of the repeat-induced overlaps, the assembler can fully resolve these repetitive regions and merge the corresponding contigs together which results in multiplicity closer to one, assembly size closer to the reference size, and reduced number of contigs. In conclusion, comparatively to the standard de novo assembly pipeline, removing 25%, 50%, and 75% of repeat-induced overlaps produces more contigs. This means even removing a subset of repeat-induced overlaps accurately, without false positives, can improve de novo assembly performance.

**Table 3:**
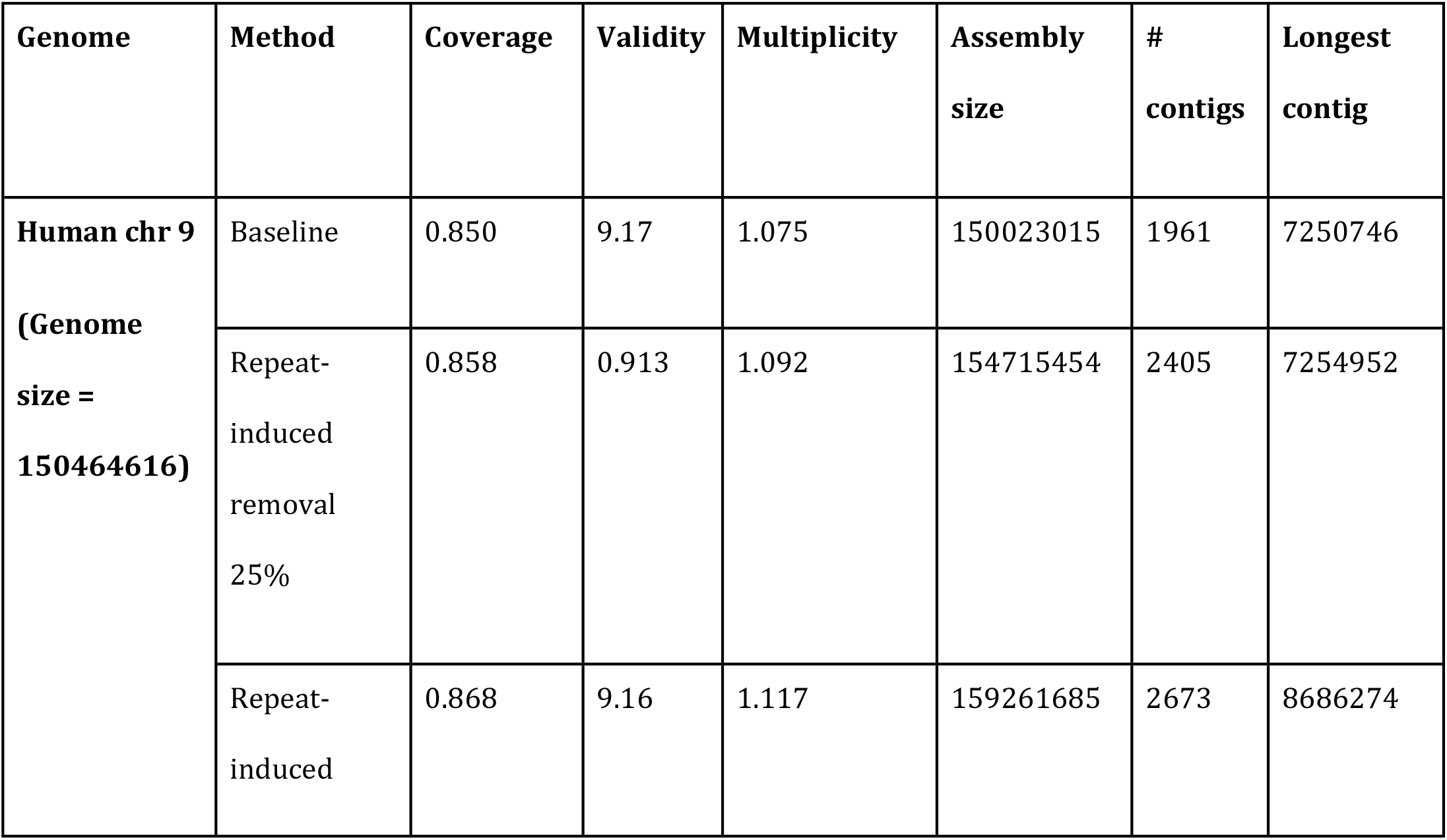
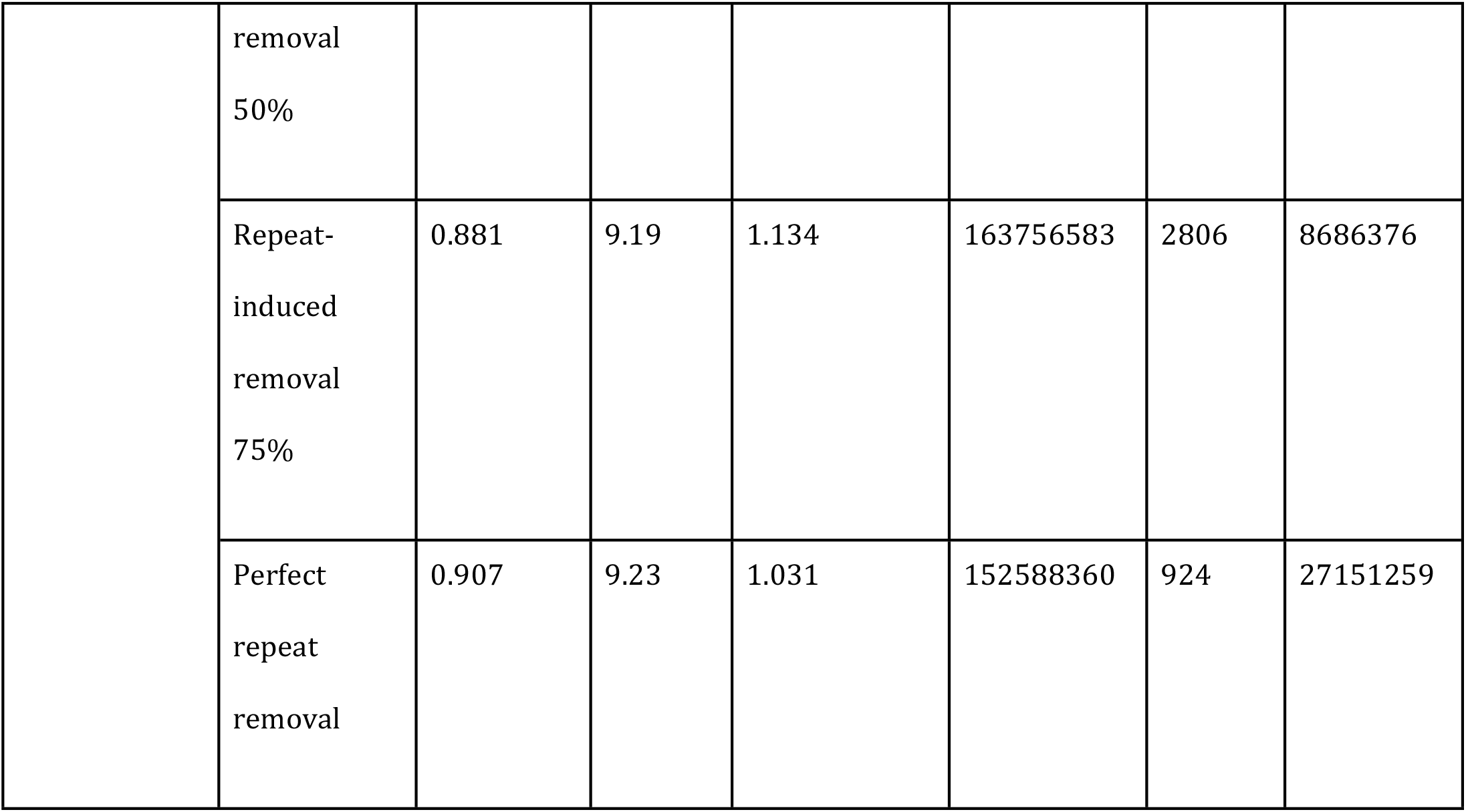
The performance of standard de novo assembly pipeline compared to de novo assembly after removing 25%, 50%, 75% and all of the repeat-induced. These metrics are described in Supplementary Table 1. With removing more repeat-induced overlaps, the coverage of assemblies is increasing. However, with removing 25%, 50%, and 75% of the repeat-induced overlaps, the number of contigs, the assembly size and the multiplicity is increasing. Meanwhile, with removing all of the repeat-induced overlaps, the number of contigs drops significantly which shows the importance of removing all of the repeat-induced overlaps.

Finally, we examined the sequence differences we got from removing the repeat-induced edges compared to following the normal genome assembly pipeline. The assembly with all repeat induced edges removed is covering additional 9476429 bp of the reference genome that is not covered in the baseline assembly. Of this additional sequence, 92% turns out to be interspersed repeat sequences. Conversely, the assembly with all repeat induced edges removed is also missing 3293397 bp with respect to the baseline assembly. Again, 93% of these are from the interspersed repeat regions. In conclusion, the majority of the newly discovered regions as well as those lost when repeat-induced overlaps were removed come from repetitive regions of human chromosome 9. It appears that repeat-induced overlaps are occasionally helpful in assembling repetitive regions, but that removing repeat-induced overlaps will result in the assembly of more repetitive regions overall.

### Training a classifier to remove repeat-induced overlaps

Since the sequence of the interspersed repeats is almost identical, we relied only on graph-based features to find and remove them. One of graph based features that can be informative to detect repeat-induced overlaps is degree. We expect the edges in the assembly graph representing repeat-induced overlaps to have a high degree since they connect two reads from the repetitive regions and those reads also align to reads originating from all other instances of the repeat. Figure 3 compares the degree of repeat-induced and normal edges in the assembly graphs. Based on Figure 3, the number of repeat-induced edges with a degree greater than 1000 is more than normal edges. However, considering edges with a degree greater than 10000, the difference is much higher, and the number of repeat-induced edges is significantly more. Therefore, we intervened in the de novo assembly process and removed the nodes representing overlaps with a degree greater than 10000 to see if removing them can improve the final assembly result. Table 4 shows the result of removing repeat-induced overlaps based on degree. No improvements are observed using this method over standard assembly pipelines. Since the yeast assembly graph does not have any edge with degree greater than 10000, we did not apply this method on it.

**Table 4:** The standard de novo assembly pipeline performance compared to perfect repeat-induced overlap removal and various repeat-induced overlap detection methods. The metrics are described in Supplementary Table 1. In all of the three test organisms, removing all of the repeat-induced overlaps improve the performance significantly, compared to the baseline scenario. In the degree method, edges with degree greater than 10000 are removed from the assembly graphs. Since the yeast assembly graph has no edge with a degree greater than 10000, we cannot apply the degree method to the yeast dataset. On the other hand, training and testing the machine-learning models require huge memory and is not achievable on the potato dataset. Our results show that, unlike the perfect repeat-induced removal scenario, these methods cannot improve the standard de novo assembly pipeline. The machine learning method results in fewer contigs compared to the standard de novo assembly pipeline, while it is losing some coverage.

| Organism | Model | Coverage | Validity | Multiplicity | Assembly size | # contigs | Longest contig |
| --- | --- | --- | --- | --- | --- | --- | --- |
| Yeast | Baseline | 0.973 | 0.943 | 1.014 | 12726687 | 33 | 958030 |
| **(Genome size = 12144833)** | Machine-learning | 0.933 | 0.934 | 1.004 | 12174134 | 29 | 1297877 |
|  | Perfect repeat removal | 0.961 | 0.934 | 1.003 | 12531324 | 25 | 1162078 |
| **Human chr 9**<br>**(Genome size = 150617247)** | Baseline | 0.811 | 0.878 | 1.082 | 150646955 | 2143 | 6179208 |
|  | Degree-based removal | 0.811 | 0.879 | 1.084 | 150834800 | 2173 | 5430347 |
|  | Machine-learning removal | 0.691 | 0.939 | 1.006 | 111503649 | 722 | 2659552 |
|  | Perfect repeat removal | 0.907 | 9.23 | 1.031 | 152588360 | 924 | 2715125<br>9 |
| **Potato**<br>**(Genome size = 731207187)** | Baseline | 0.631 | 0.945 | 1.068 | 522035794 | 12794 | 315461 |
|  | Degree-based removal | 0.629 | 0.945 | 1.069 | 520480215 | 12796 | 315461 |
|  | Perfect repeat removal | 0.701 | 0.941 | 1.008 | 549511126 | 11805 | 315508 |

Another way to detect repeat-induced overlaps is to train a machine learning-based classifier based on graph-based embedding. First, we generated separate train and test datasets to evaluate this method fairly. We simulated two reference sequences based on the reference genome of the three organisms we analyze. After that, we simulated reads from these simulated reference sequences and performed a pairwise alignment between the reads. We used the reference genome and the first simulated read set to train and test the GraphSage embedding model. To train the GraphSage embedding, we select subgraphs using StellarGraph’s edgesplitter method. Then we labeled each pair of nodes in the subgraph as 0, 1, 2 where 0 represents normal edge, 1 repeat-induced edge, and 2 no edge. Table 5 shows the performance of the GraphSage embedding model on train and validation data. Interestingly, the model is not efficient in separating the three classes of edges in the yeast dataset, while it is performing well on human chromosome 9 dataset.

**Table 5:** This table shows the performance of the GraphSage embedding model and the logistic regression classifier. We use the edgesplitter module in the StellarGraph library to sample subgraphs for the train and test datasets. The size of subgraphs is 20% and 2% of the actual yeast’s and human’s assembly graphs, respectively. To test the performance of the logistic regression classifier, we use a 10-fold cross-validation. Interestingly, the human GraphSage and logistic regression models perform better than the yeast ones, showing more significant differences between the repeat-induced and normal edges in the human assembly graph.

| GraphSage model |  |  |  |  |
| --- | --- | --- | --- | --- |
| Metric | Train accuracy |  | Validation accuracy |  |
| Yeast | 0.5356 |  | 0.5387 |  |
| Human chromosome 9 | 0.7653 |  | 0.7646 |  |
| Logistic regression classifier |  |  |  |  |
| Metric | F1 score (SD) | Accuracy (SD) | Precision (SD) | Recall (SD) |
| Yeast | 0.761 (0.007) | 0.936 (0.002) | 0.788 (0.008) | 0.740 (0.007) |
| Human chromosome 9 | 0.887 (0.001) | 0.911 (0.001) | 0.915 (0.001) | 0.868 (0.001) |

Next, we used the extracted embeddings of overlaps in the second simulated dataset to train a classifier for separating normal and repeat-induced overlaps. Since the dataset is imbalanced, and the graphs have more normal edges in yeast genome and more repeat-induced edges in human, we up-sampled and down-sampled repeat-induced edges in yeast and human datasets, respectively. Following that, we trained a logistic regression classifier and evaluated it with 10-fold cross-validation (see Table 5). While the GraphSage embedding model failed to separate the three classes of edges in the yeast dataset, the logistic regression classifier achieved impressive results in separating repeat-induced and normal edges using the same embedding model on the second simulated dataset. Interestingly, the GraphSage model performed much better on the human chromosome 9 assembly graph and achieved 76% validation accuracy.

Last, we extracted the embeddings of overlaps in the last dataset and used the classifier trained in the previous step that achieved the highest F1 score to predict the repeat-induced overlaps. After removing the overlaps predicted as repeat-induced, we assembled the remaining overlaps and evaluated the results (see Table 4). The performance of yeast assembly drops after removing the overlaps predicted as repeat-induced. That means that the disadvantage of losing some of the normal edges in the yeast assembly graph because of prediction errors is more than the advantage of removing repeat-induced overlaps. Since the yeast genome does not have many interspersed repeats and repeat-induced edges (see Tables 1 and 2), this is not surprising. On top of that, the only feature we assigned to the nodes before training the GraphSage model is the degree of nodes, while in the yeast assembly graph, the degree of repeat-induced and normal edges is not significantly different (see Figure 3.a). However, the length of the longest contig is increased, and the number of contigs is reduced, which shows that the method solved the previously challenging repetitive regions.

Similar to yeast, human chromosome 9 assembly performance is lower than baseline after removing overlaps predicted to be repeat-induced (see Table 4). The coverage is ∼12% lower and the assembly size is ∼40Mbp smaller than the actual chromosome 9 size. The number of contigs is smaller than all the other cases, and the multiplicity and validity are close to one, which means the assembly and reference map are nearly one-to-one. As a result, the machine learning method is successful in removing some essential repeat-induced overlaps, which enables the assembler to merge the contigs that were split apart before. However, the model also incorrectly predicts some critical normal overlaps as repeat-induced, resulting in decreased coverage and assembly size when they are removed. Despite our best efforts, we were unable to apply the machine-learning method to the potato dataset due to its large size and memory requirement.

## Conclusion

In this study, we study the effect of interspersed repeats on de novo genome assemblies of three organisms, i.e., yeast, human chromosome 9, and potato. The reads originating from interspersed repeat regions align with those from all instances. Therefore, it is possible to label the alignments with not overlapping originating coordinates as repeat-induced overlaps. Here, we analyze the effect of repeat-induced overlaps in the assembly graph and de novo assembly. At last, we investigate some strategies to detect and remove repeat-induced overlaps.

Interspersed repeats make up approximately 1, 6, and 10% of the yeast, human chromosome 9, and potato genomes, respectively. Although the repeats are causing only 1% of the overlaps in the yeast dataset, they correspond to 76% and 96% % of overlaps in human and potato datasets. Since most of the overlaps in the assembly graph of these two genomes are repeat-induced, this is the most challenging problem to solve in genome assembly.

To investigate the effect of repeat-induced edges in the assembly graph on the final assembly result, we removed all of the repeat-induced overlaps and compared the results to the normal de novo assembly pipeline. We observed that removing repeat-induced overlaps improved coverage and continuity of the assembly, even in yeast with much lower repetitive content. In potato, which has the most repetitive contents among the test organisms, removing repeat-induced edges leads to a 9% improvement in coverage.

We investigate if it is possible to detect repeat-induced overlaps based on the degree of their corresponding edges in the assembly graph. We define the degree of an edge as the sum of the degree of two nodes connecting the edge. As shown in Figure 3, most of the repeat-induced overlaps in human chromosome 9 and potato assembly graphs have more than degree 10000. Therefore, we remove edges with more than degree 10000 and see the effect of it on the final assemblies. As shown in Table 4, there is no improvement in the assemblies after removing edges with degree greater than 10000, and the final assemblies are very close to the standard assembly pipeline.

We also attempt to train a classifier to detect repeat-induced edges based on graph-based features. Although we achieved some improvement after removing repeat-induced edges with the classifier, the results are far from the results when all of the repeat-induced edges are removed. This shows great potential for a follow-up project to detect and remove repeat-induced overlaps accurately.

We suggest that detecting and removing repeat-induced overlaps can be one a smart edge filtering method during assembly. Our attempt to train a classifier that accurately detects and removes repeat-induced overlaps did not achieve significant results. However, our results show that a perfect classifier that removes all the repeat-induced overlaps can make impressive improvements in the genome assembly process.

